# A scaling law relating the rate of destruction of a solid tumor and the fractal dimension of its boundary

**DOI:** 10.1101/2022.06.29.498072

**Authors:** Álvaro G. López, Lorena R. Sanjuán

## Abstract

We investigate the scaling law relating the size of the boundary of a solid tumor and the rate at which it is lysed by a cell population of non-infiltrating cytotoxic lymphocytes. We do it in the context of enzyme kinetics through geometrical, analytical and numerical arguments. Following the Koch island fractal model, a scale-dependent function that describes the constant rate of the decay process and the fractal dimension is obtained in the first place. Then, *in silico* experiments are accomplished by means of a stochastic hybrid cellular automaton model. This model is used to grow several tumors with varying morphology and to test the power decay law when the cell-mediated immune response is effective, confirming its validity.

## 1. Introduction

Along the last decades, the usage of monoclonal antibodies to modulate T-cell immune checkpoint molecules, such as CTL4-activity and the blocking of the receptor PD1 with the anti-PD1 antibody, have impaired a renewed impetus to the study of immunotherapy in the fight against some cancers such as advanced melanoma (Waldman et al., 2020). Given the complex regulatory signaling pathways that control both cell growth and the cell-mediated immune system response, our capacity to understand their molecular and cellular basis partly relies on our ability to develop mathematical models (Wodarz and Komarova, 2014). These models might be indispensable to explain intricate dynamical aspects of such molecular and cellular mechanisms. Theoretically, they also provide an analytical framework where fundamental questions about cancer dynamics and evolution can be addressed in a more rigorous way. Ultimately, their development is of practical importance, since they might permit the improvement of the existing therapies, or even suggest new strategies (Kirschner and Panetta, 1988; López et al., 2017b).

Many mathematical works on the modeling of the interactions between tumor cells and cytotoxic effector T-cells have shown that their dynamics can be described in terms of modified enzymatic reactions (Hill, 1910; Monod, 1949; Johnson and Goody, 2011), where the tumor cells play the role of the substrate and the immune cells play the role of the enzyme (see Fig. 1). Originally proposed in the mid 90’s (Kuznetsov et al., 1994), this idea has been developed in mathematical models of tumor-immune interactions (De Pillis et al., 2005; López et al., 2014) validated against data gathered from chromium release assays, *in vivo* experimental results (Diefenbach et al., 2001; Dudley et al., 2002), and also more recently using *in silico* experiments with stochastic hybrid cellular automata models (López et al., 2017a). Models inspired by the Michaelis-Menten kinetics are frequently used to represent both immune cell recruitment and tumor cell lysis (Kirschner and Panetta, 1988; Kuznetsov et al., 1994; De Pillis et al., 2005). In the present work we shall focus on the process of tumor lysis in immunocompetent environments.

**Figure 1:**
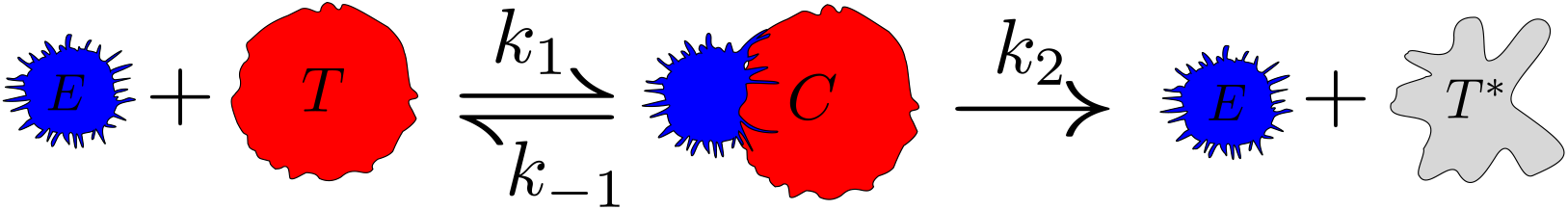
Cytotoxic T-cell mediated immune response. An activated lymphocyte *E* couples to a tumor cell *T*. After receptor tumor cell recognition by the immune cell, the two cells couple forming a complex *C*. The result of the interaction is the initial immune cell and an apoptotic tumor cell *T*\*. The whole process of lysis frequently lasts around forty minutes (Weinberg, 2006). This cellular interaction is similar to an enzymatic chemical reaction, where the tumor cell plays the role of the substrate and the immune cell acts as an enzyme.

The idea of using chemical reactions as a metalanguage to model dynamical processes at higher scales of the biological hierarchy is very old. It can be traced back to mathematical population models, originally developed by Leonardo de Pisa, Leonhard Euler and Francoise Verlhust (Bacaër, 2011). This tradition was continued by many other important authors, and has been epitomized by the predator-prey ecological model devised by Alfred Lotka (Lotka, 1920), which was also used by Vito Volterra to describe fish catches (Volterra, 1926) in the Adriatic Sea. However, all these approaches present noticeable difficulties. Among them, perhaps the most outstanding one is the fact that the *law of mass action* is not expected to hold when the size of the interacting elements is relevant to the scale at investigation. At the macroscopic scale, systems are not diluted let alone well stirred, not even locally. Therefore, we can not expect the chemical law of mass action to rigorously hold. For example, when the individuals of two interacting animal species come together (collide, if desired), they usually keep engaged for a while and reaction proceeds with great probability. Consequently, saturation effects limiting the speed of the reaction can appear rather soon during dynamical evolution. In this regard, the simple incorporation of spatial dependence to the evolving population densities, as it is frequently done in order to develop reaction-diffusion models, is also subject to the same hesitations. In all these situations the spatial limitations due to the geometrical arrangement of the interacting elements becomes unavoidable. These considerations appear sometimes in more sophisticated ecological laws beyond the law of mass action, such as in the Holling *functional responses* describing interacting animal populations in ecology (Holling, 1959), or as in the Arditi-Ginzburg equation (Arditi and Ginzburg, 1989), where ratio-dependent rates are considered. Just to recall, in these models the pray plays the role of the substrate, while the predator represents the enzyme.

In order to accommodate the Michaelis-Menten kinetics to the tissue scale when tumor-immune interactions are studied, a key modification has been recently proposed (López et al., 2017a). This proposal makes the constant rates of the reaction depend as power laws on the enzyme and the substrate concentrations. In this way, the production of products is slowed down as saturation of the immune cell concentration tends to increase way beyond the tumor cell concentration. We recall that the conventional Michaelis-Menten kinetics already incorporates saturation due to the substrate, but does not cover cases in which the enzyme concentration saturates. Indeed, these two processes are not entirely symmetric, for an enzyme can catalyze several or even many substrate molecules before inactivating, whereas products rarely return to their substrate configuration after the interaction. Be that as it may, this approach proves that sometimes it is possible to incorporate the effects of spatiotemporal phenomena related to very heterogeneous reaction-diffusion processes indirectly in ordinary differential equation models, without appealing to complicated spatiotemporal discrete models.

None of the previous works has provided a strict analytical mathematical function relating the fractal dimension of the boundary of the tumor (here playing the role of the prey) and the maximum rate of tumor cell depletion, when the effector-to-target ratio is sufficiently high. The importance of fractal dimension in solid tumor growth is well documented and has proved of great assistance to diagnose and treat some malignancies (Baish and Jain, 2000; Norton, 2005; Goh, 2009). In the present work we analytically derive this scaling relation and test its accuracy by means of a cellular automaton model.

## 2. The kinetics of tumor lysis

The differential equation (López et al., 2016; López et al., 2017a) describing the velocity at which a population of tumor cells *T* integrating a compact solid tumor, is destroyed by the cell-mediated arm of the immune system, can be written as 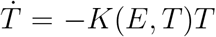, with *K* the fractional cell kill function, which is defined as *K*(*E,T*) = *rE*^*λ*^*T*^*ν*−1^*/*(*sT*^*ν*^ + *E*^*λ*^). The fractional cell kill describes the rate at which the immune cell population of effector cells *E* reduces the tumor to a certain fraction of itself. Assuming that the tumor cells do not reproduce, as if they had been irradiated, the velocity at which a tumor is lysed is governed by the differential equation

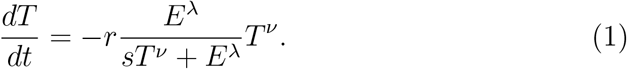

For a fixed value of effector cells *E*, this equation can be solved in terms of ordinary hypergeometric functions. The parameter *r* represents the maximum constant rate at which the tumor cell population decays, when the immune cell population is overwhelmingly high in comparison to the tumor (*E*^*λ*^*/sT*^*ν*^ → ∞). It has been shown that less spherical tumors lead to higher values of this parameter (López et al., 2016; López et al., 2017a). The parameter *ν* is given by the topology of the tumor in relation to the immune cell population. It takes a value *ν* = 1 − 1*/D* for connected solid tumors non-infiltrated with lymphocytes, where *D* is the topological dimension of the space in which the tumor is embedded. On the opposite extreme, the constant *ν* increases up to a maximum value of one as the infiltration is more pronounced or when the malignancies are of haematological origin (López et al., 2016). In this extreme case the decay of the tumor is exponential when the immune cells completely infiltrate it, as it occurs with radioactive decays. The parameter *λ* depends on the relative spatial disposition and contact between the tumor and the immune cell population. Finally, the parameter *s* is related to the intrinsic ability of the cytotoxic cells to recognize and destroy their adversaries (López et al., 2017a). Smaller values of this parameter are associated with a more efficient immune system.

The kinetic law represented by Eq. (1) is stated as follows. The velocity at which a tumor is lysed by an effector cell population becomes faster as this immune cell population increases. Once the immediate surroundings of the tumor domain are profusely occupied by some layers of immune cells, saturation is attained due to *cell crowding*, because the rest of the immune cells are not in contact with their tumor adversaries. Naturally, this saturation process depends on the size and geometry of the tumor. On the other side, when the tumor cell population increases from zero (immune cell numbers kept fixed), faster lytic velocity is obtained, but saturation befalls once again. The reason is that for a too big tumor compared to the immune cell population, the addition of tumor cells can barely increase the rate at which the tumor is lysed, since these added tumor cells are not in contact with immune cells, which are already busy lysing some of the initial tumor cells. Certainly, it takes some time for a CTL to destroy a tumor cell, and even if this time is small, it is well known that immune cells inactivate after several encounters with tumor cells (Kuznetsov et al., 1994).

We now briefly examine the two limits appearing in this equation. For a fixed number of immune cells *E*, when the immune cell population is small compared to the tumor size (*E*^*λ*^ ≪ *sT*^*ν*^), the tumor cell population is reduced at a constant velocity 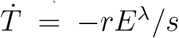. This uniform decay represents the extreme situation in which a few lymphocytes are fighting a comparatively big tumor. Ideally, if it takes these immune cells around forty minutes to lyse a tumor cell (Weinberg, 2006), the expected velocity of the decay is simply a few tumor cells per hour. Of course, the rate of destruction clearly depends on the intrinsic ability of the cytotoxic cells *s* to lyse the tumor cells, and also on the tumor morphology, which is coded in the parameters *r* and *λ*.

The second limit occurs when the immune cell population is high enough compared to the tumor cell population (*E*^*λ*^ ≫ *sT*^*ν*^). Then, the Eq. (2) yields an *algebraic decay*, which for two-dimensional tumors is described by the equation

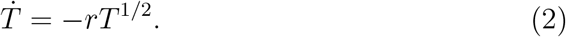

This other scenario corresponds to the case in which the tumor surroundings are totally covered with effector cells. Interestingly, when combined with the growth of the tumor, this power law can lead to new unexplored *non-analytic bifurcations*, different from those appearing in the normal forms of traditional bifurcation theory (López et al., 2017c). The solution to this equation leads to a parabolic decay of the tumor. A recently published theorem (López et al., 2017c) proves that, in this limit, the constant rate *r* of the tumor can be written as

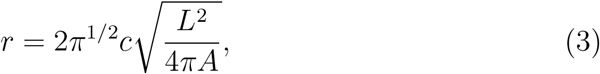

where *c* is the rate at which one immune cell destroys a single tumor cell, *L* is the perimeter of the tumor, and *A* represents its area at the time when the lysis process starts *t* = 0. Thus in the absence of other phenomena, the isoperimetric inequality ensures that spherical tumors protect better themselves from immune attacks (Osserman, 1978). It is for this reason that the term inside the square root in Eq. (3) has been called the *sphericity*, ranging from zero to one (Wadell, 1935). A similar equation can be derived for three-dimensional tumors, to which the present analysis can be perfectly extended. The total time *t*_*tot*_ it takes a tumor to be fully erased, according to Eq. (2), can be estimated as

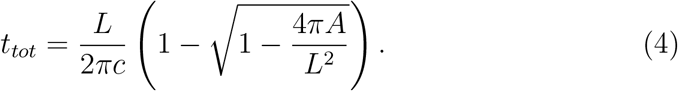

Again, the reader can check that spherical tumors display the maximum total time *t*_*tot*_ = *L/*2*πc*. In the following lines we show that the length and the area of the boundary scale as a nontrivial function of the fractal dimension of the tumor, which in the appropriate limit approximates to an ordinary power law. As it is well known, this behavior is typical of fractal geometrical objects, which are ubiquitous in biology.

## 3. The scaling law

We now obtain a relation between the length *L* of the perimeter of the tumor, and also the area *A* that it encloses, as a function of the scale *ε*. This parameter reveals the fine structure of the tumor’s boundary as one zooms in, after having settled at some point of such a boundary. Two extreme scales must be considered when studying a tumor: the one comprised by the tumor as a whole *ε*_0_, and the one *ε* determined by the elements that constitute the tumor, *i.e*, the eukariotyc human cell. To put some numbers to these magnitudes, if we consider that the radius of a typical cell rounds *ε* = 10*μm*, and that a detectable three-dimensional tumor of one gram comprises around 10^9^ cells, we can estimate that *ε*_0_ = *L*_0_ = 10*mm*. This yields a ratio *ε*_0_*/ε* of three orders of magnitude.

Just to derive the mathematical functions *L*(*ε*) and *A*(*ε*), we instrumentally use as a guiding model of a tumor the Koch island (Koch, 1904). As depicted in Fig. (2), if the edge of the starting triangle has a size of *L*_0_, then the length of the curve at the *n*-th iteration is *L*_*n*_ = 3*L*_0_ · (4*/*3)^*n*^, which diverges to infinity as the fine structure of the curve reveals. The scale reduces as *ε*_*n*_ = *L*_0_(1*/*3)^*n*^. The homothety dimension of the Koch island is *D*_*f*_ = log 4*/* log 3. For perfectly self-similar geometrical objects this dimension coincides with the more general box-counting dimension, which is computed as

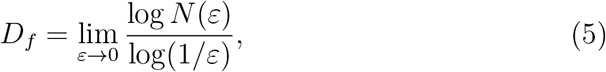

where *N* (*ε*) is the number of edges required to close the Koch curve at the scale *ε*. By replacing *ε* with *ε*_*n*_ we find that 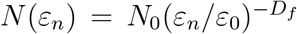, and obtain immediately the result for the Koch curve. Thus, in general, we can write the scaling law 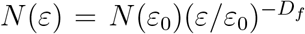. This entails us to write the perimeter length scale relation for a general fractal curve as 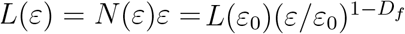.

The scaling law of the area is not much harder to obtain. As shown in Fig. 2, for the Koch island the area is obtained by adding triangles whose area scales as 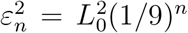. In the first step, we add three triangles, in the second 3 · 4 triangles, and in the *n*-th step 3 · 4^*n*^ triangles are added.

**Figure 2:**
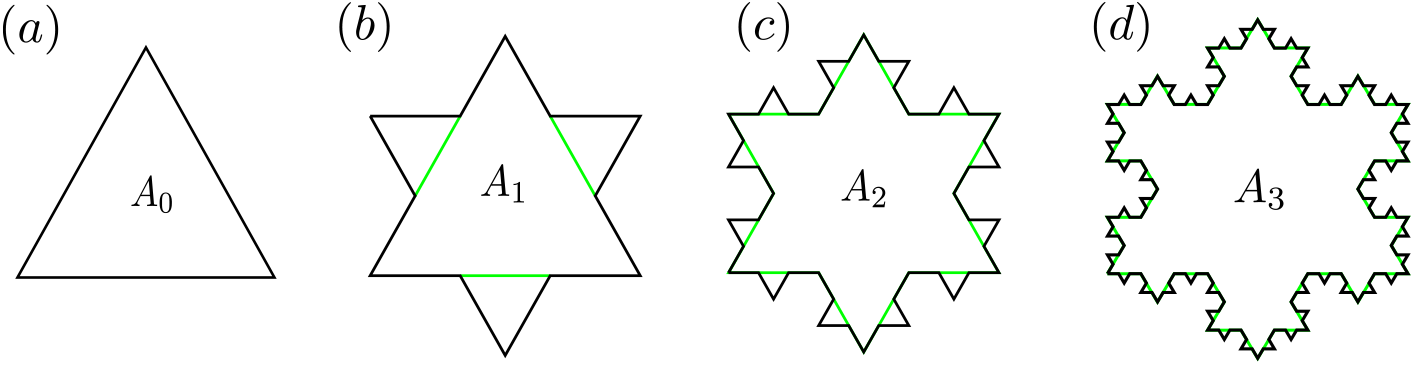
Koch island. The first four iterations of the construction of the Koch island, starting with a triangle that is scaled by one third of its original size, cloned three times and glued to the middle segments of the edges of the initial triangle. If we repeatedly iterate this process, a compact set whose boundary has infinite perimeter length and that is everywhere non-differentiable, is generated. The total area for each iteration *A*_*k*_ has been remarked, which grows monotonically and converges to a finite value. It constitutes the main interest of the present fractal model.

Therefore, the total area at the *n*-th iteration can be computed as the sum of the areas of all the triangles, leading to the series

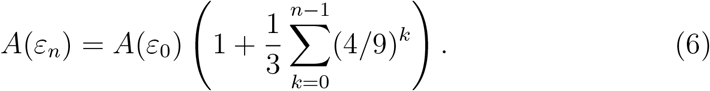

If we note that the 1*/*3 factor is determined by the initial geometrical figure, and also that *N* (*ε*_*k*_)*/N*_0_ = 4^*k*^ for the fractal Koch curve, we can write the following general formula for the area of a two-dimensional simply connected and compact fractal object

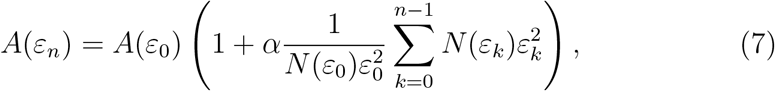

where *α* is a factor that depends on the initial starting set. By considering that 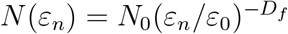, we obtain

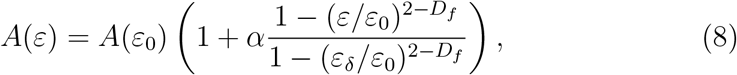

where *ε*_*δ*_*/ε*_0_ is the value of the homothety that is operated at each step. For the Koch curve we have a value 1*/*3. In the limit when *ε* tends to zero, Eq. (8) gives the area of the Koch island. Replacing these values in Eq. (3) allows us to obtain a scaling law between the cell-to-tumor size ratio and the fractal dimension of the boundary as

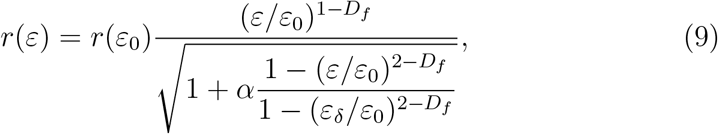

with *r*(*ε*_0_) given by Eq. (3), but at the original scale. In the limit in which 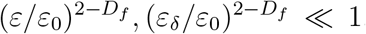, which holds reasonably well for a detectable solid tumor of fractal dimension not too close to two, we can approximate the power law as

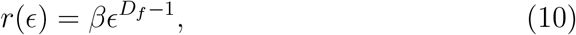

where the renormalized scaling parameter *ϵ* = *ε*_0_*/ε* and the parameter 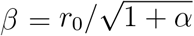 have been introduced for simplicity. This new analytical result predicts a linear relation between the log *r* and the fractal dimension of the boundary *D*_*f*_ of the tumor. It is in conformity with claims made in previous works (López et al., 2016; López et al., 2017a). Namely, *as the fractal dimension of the boundary of the tumor increases, the rate at which it is predated by the immune system increases as well*, ceteris paribus. More generally stated, spherical arrangements protect better their interior from external attacks, by reducing surface contact. Of course, for this precise reason, this protection has its disadvantages too: spherical tumors mostly grow from their boundary and frequently display necrotic cores at their centre due to a lack of nutrients and oxygen, and because of acidification (Bru et al., 2003; Chandrasekaran and King, 2012). In the forthcoming sections we present the computational tools needed to test the accuracy of this power law and perform such tests.

## 4. A hybrid cellular automaton model

A stochastic hybrid cellular automaton (CA) model designed to study the growth of avascular tumors has been used in previous works (Ferreira et al., 2002; Mallet and De Pillis, 2006; López et al., 2016; López et al., 2017a, 2019). In Fig. 3 a schematic representation of this model is depicted. We refer the reader to these works for a rigorous inspection of the detailed CA rules, together with the equations for the diffusion of nutrients (*e.g*. glucose and oxygen) from the capillaries to the tissue environment (López et al., 2017a). Here the process of lysis has been considerably simplified, as compared to previous models. We summarize the most relevant features concerning the growth of the tumors, and then proceed to thoroughly explain the adaptations that we have performed to test the decay law appearing in Eq. (10).

**Figure 3:**
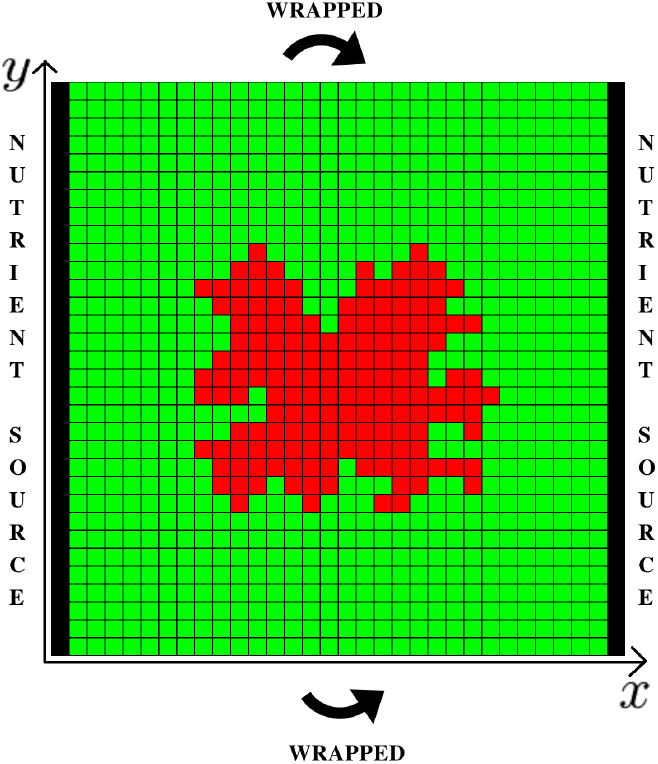
The cellular automaton. The tumor cells occupying several positions in a square grid are shown in red. The other spots are occupied by the healthy cells. The black columns in the left and right boundaries of the square represent the vessels from which nutrients diffuse. Periodic boundary conditions are assumed in the upper and the lower sides of the domain.

**Figure 4:**
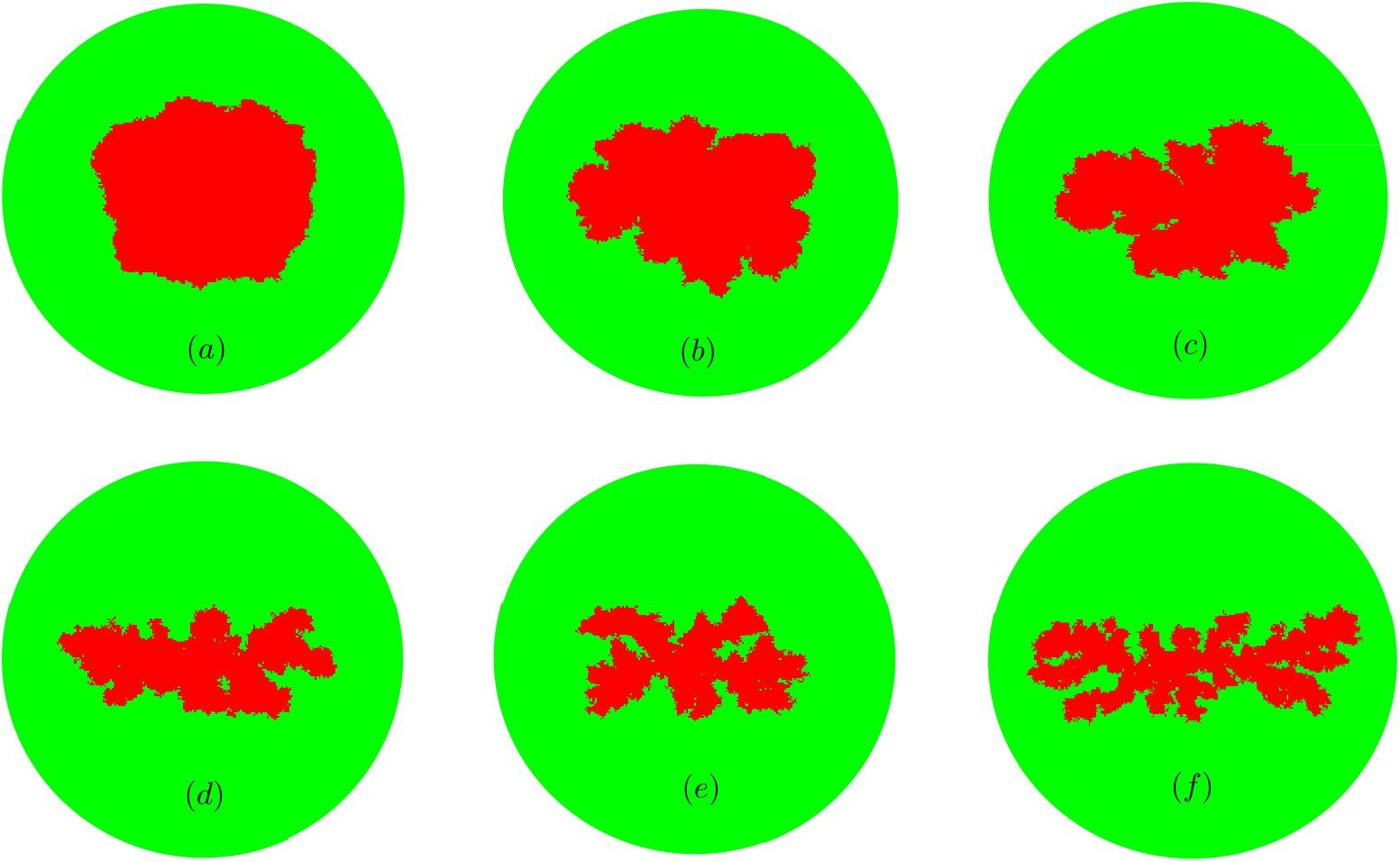
Six tumors grown by iteration of the cellular automaton. A grid of *n* × *n* cells, with *n* = 300 has been used. We disregard necrosis and motility of tumor cells by setting the parameters *θ*_*div*_ = 0.3, *θ*_*nec*_ = 0 and *θ*_*mig*_ = ∞. In all the six cases *λ*_*M*_ = 10. (a) A spherical tumor obtained for parameter values *α* = 0.5*/n* and *λ*_*N*_ = 6. (b) A spherical to papillary tumor is obtained for parameter values *α* = 3.3*/n, λ*_*N*_ = 140. (C) A papillary tumor obtained for parameter values *α* = 4.1*/n* and *λ*_*N*_ = 190. (d) A papillary to filamentary tumor obtained for parameter values *α* = 5.9*/n* and *λ*_*N*_ = 222. (e) Another papillary but more filamentary tumor with the parameter choice *α* = 6.8*/n* and *λ*_*N*_ = 232. (f) A filamentary tumor with the parameter choice *α* = 8*/n* and *λ*_*N*_ = 270. These six tumors have grown up to numbers between 9000 to 10000 cells.

### 4.1. Tumor growth

As in any hybrid CA model, we represent the cells discretely, allowing them to occupy any of the grid points of a square lattice, representing the spatial domain. The *in silico* tumor cell cultures are comprised by two different types of cells: healthy cells and tumor cells. On the other side, the nutrients diffuse from the vessels into the domain, and they are described through linear reaction-diffusion equations, which are continuous and deterministic. The dynamical evolution of cells and nutrients is governed by the following rules, respectively:

1. Each type of action *a* of the CA cells occurs according to probabilistic rules depending on some parameters *θ*_*a*_ and the nutrient concentration at each point of the grid. These parameters represent the intrinsic ability of the cells to carry out the corresponding action. The role of healthy cells is passive, consuming nutrients and being replaced by tumor cells. Tumor cells can reproduce *θ*_*div*_, migrate *θ*_*mig*_ or die *θ*_*nec*_. For the present work, where compact solid tumors not infiltrated with lymphocytes are studied, migration has been disregarded.
2. We distinguish two types of nutrients: those that are specific for cell division *N* and those that are related to the remaining cellular activities *M*. The rate of nutrient consumption is represented by the parameters *λ*_*M*_, *λ*_*N*_ and *α*, respectively. On the vertical sides of the domain we impose Dirichlet boundary conditions, right where the vessels are placed. Periodic boundary conditions are imposed in the upper and the lower sides of the square.

### 4.2. Tumor lysis

During the second step, we peel the tumors layer by layer at each step in a completely deterministic way. Each cell of the tumor’s boundary is erased at each step and then replaced by an immune cell. In other words, immune cells lyse with probability value of one, and recruit also surely, so that the tumor is completely covered with immune cells at all times. This simplification speeds enormously the algorithm and, as has been shown in previous works (López et al., 2016), yields the same results as compared to the conventional CA probabilistic rules, in the limit in which the immune cells are placed isotropically covering the whole tumor and are thus renewed after inactivation through the mechanism of T-cell recruitment.

### 4.3. The algorithm

As in previous works (De Pillis et al., 2005; López et al., 2014; López et al., 2016; López et al., 2017a), which are inspired by chromium release assays, the simulations are conducted in two steps. Six tumors with varying geometry are grown in the first place. Then, the healthy tissue surrounding each tumor is numerically washed out and replaced by immune cells. Because we are only interested in understanding the dynamics of the process of lysis when the immune response is strong, we neglect the growth of the tumors during the action of the immune system, as if they had been irradiated. A complete study covering the transient and asymptotic dynamics of tumor-immune aggregates has been provided somewhere else (López et al., 2017d). We also assume that the immune cells advance synchronously, replace the destroyed tumor cells and never inactivate. Otherwise, if the immune cells inactivated, they would be quickly replaced by other immune cells which advance towards the tumor or by the abovementioned process of immune cell recruitment (López et al., 2017d). These two considerations allow us to simplify noticeably the algorithm, by neglecting T-cell recruitment, their migration and the inactivation of T-cells. Summarizing, we just erase each outermost layer of the tumor step by step.

## 5. Results

The results of the CA simulations are shown in Fig. 6. A total sample of six tumors with increasingly fractal morphology have been generated using our cellular automaton, ranging from spherical-like geometries to papillary tumors and, finally, to filamentary tumors displaying many branches. The phenomenon producing the spherical *symmetry breaking* of the tumors is the result of increasing nutrient competition between tumor cells in the model. A positive feedback establishes between growth towards the vessels and nutrient consumption. This effect is sometimes called the *Mathew effect* in other branches of science (Merton, 1968) and has been thoroughly explained in previous works (López et al., 2017a). Briefly, when a group of cells divide towards the vessels, even randomly selected, they begin to self-promote their growth consuming plenty of nutrients and leading other neighboring parts of the tumor to starvation.

Once the tumor cell cultures achieve a size close to 10^4^ cells, their growth is halted and the immune cell-mediated immune response begins, as explained in the previous section. We have assigned a time of one hour to each step of the cellular automaton hereafter, which now operates deterministically. Considering stochastic laws does not introduce noticeable differences when the immune system is efficient and covers the entire tumor, as shown in previous works (López et al., 2016). During each step, the most external layer of a tumor is completely erased by the immune cells. As can be seen in Fig. 5, all the tumors fit very well the power decay law 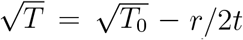, with rather unbiased residues, which solves the differential equation 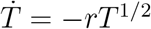. Some tendency to coherently oscillate around the deterministic law is observed, possibly attributable to the fact that the tumors tend to change their geometry as they are lysed. The last steps of the tumors’ decay have been neglected, since a description based on continuous population cell densities breaks at low cell numbers.

**Figure 5:**
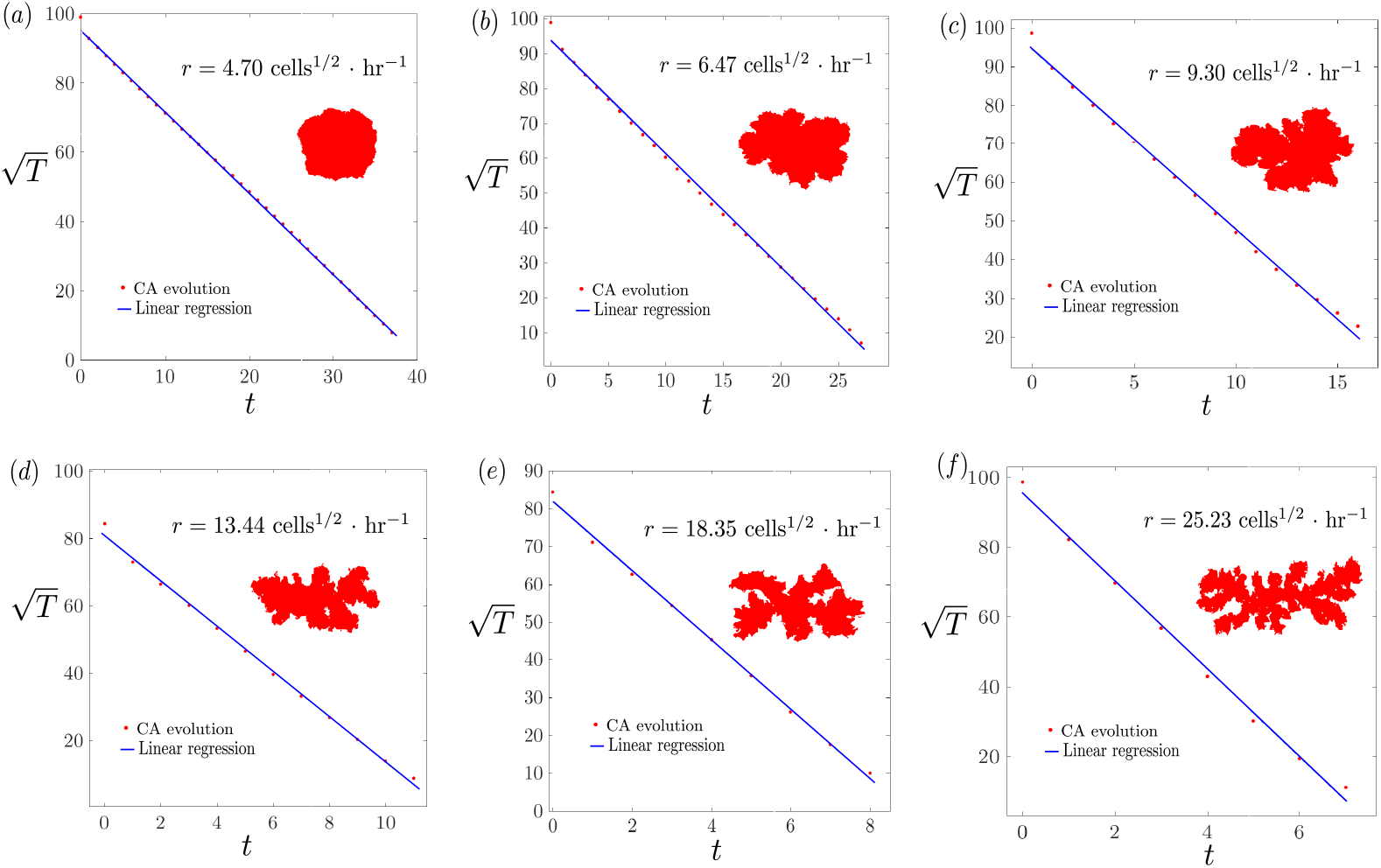
Tumor lysis. The six tumors computed with the CA are synchronously erased layer by layer, and fitted to the law 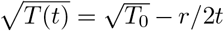. The growing value of the decay constant parameter *r* is displayed on the top right margin of each graphic. (a) A tumor with a rather spherical shape decays to negligible cell numbers in almost 40 hours. (b) A spherical to papillary tumor reduces in 26 hours. (c) A papillary tumor decays faster, in 17 hours (d-e). As the shape of the tumor turns from papillary to filamentary, the tumor can decay in 10 or 8 hours, respectively. (f) It takes the filamentary tumor to decay not much more than 6 hours, displaying the fastest decay rate.

As described in previous works (López et al., 2016), and it is also evident from Fig. 6, the parameter *r* describing the decay rate increases for higher fractal dimensions of its boundary. The fulfillment of the new scaling law appearing in Eq. (10) is inspected in Fig. 6, confirming this functional relation. Again, we distinguish randomly distributed residues, but now plotted inside the figure. Importantly, we can compute from the fitting log *r* = *D*_*f*_ log *ϵ* + *c* a value of log *ϵ* = 6.32, which yields a result quite close to the expected value for a detectable tumor of one gram, according to the estimations provided at the beginning of Sec. 3.

**Figure 6:**
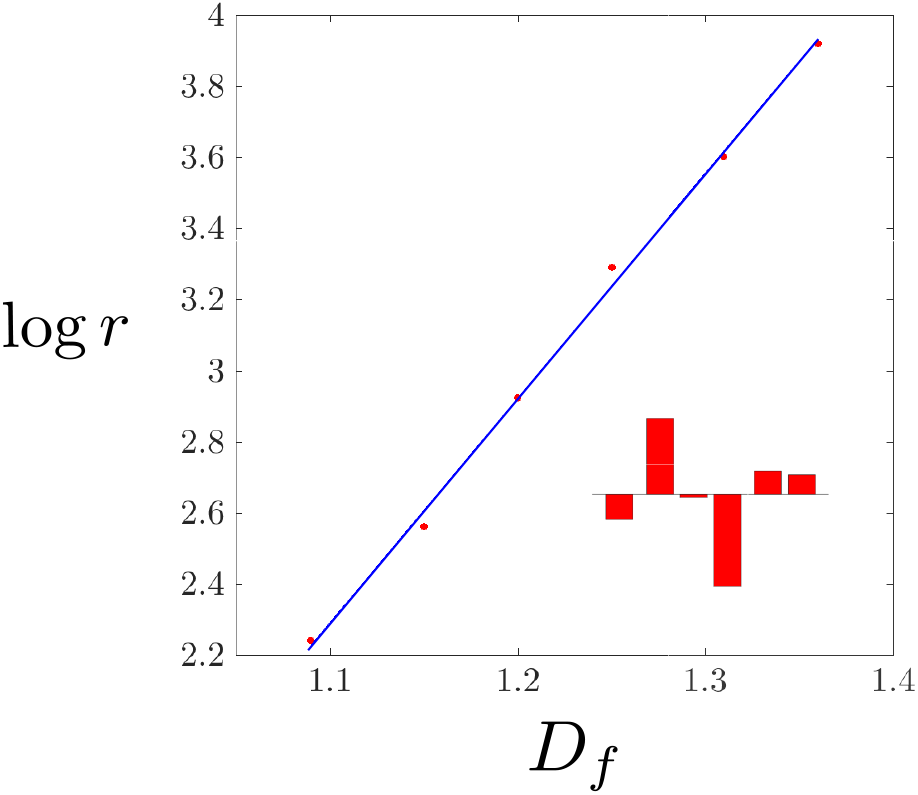
Decay rate and fractal dimension. The function 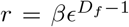 is adjusted to a straight line by a least square fitting method. The data follow nicely the scaling law, with randomly distributed residues (see the red bars for their relative size and sign), confirming its tendency.

## 6. Discussion

We have demonstrated that there is a power law relation between the fractal dimension of the boundary of a tumor and the constant rate at which it is lysed by a population of cytotoxic immune cells. The scaling law is nicely satisfied. We expect it to hold as long as the fractal dimension of the tumor boundary is rather close to one, the immune response is strong and there is not a severe degree of T cells infiltration into the tumor. Otherwise, if the immune effector cells penetrate profusely into the solid tumor, an exponential decay must be attained, according to Poisson statistics (López et al., 2016). Thanks to the present work, the status of the mathematical law represented by Eq. (10) is such that two of its parameters (*ν* and *r*) can be estimated from analytical arguments solely in terms of the tumor morphology and its topology. However, there are still two other parameters relying on the fitting of lysis curves to experimental or numerical data (López et al., 2014, 2017a).

Concerning the parameter *λ*, we do not expect it to be easily explained by analytical reasoning, since it depends on the degree of contact between the tumor and the immune cells, which one way or another must also affect the other parameters to some extent. It is worth asking if an estimation of the parameter *s*, which describes the efficiency of cytotoxic T cells, can be provided in terms of the analysis of the immune T cell and the tumor cells phenotype profiles, *e.g*., the determination of the levels of ligands expression, such as the ligands of the NKG2D receptor (Diefenbach et al., 2001). A connection between them has already been suggested through bifurcation theory (López et al., 2014), but without rigorously entering into the question of how this parameter emerges from the average levels of ligands expressed in tumor cells, and other microscopic molecular features as well.

To conclude, some remarks are deserved about the extent to which Eq. (1) can be adapted to broader macroscopic systems, including predator-prey dynamics and, more generally, enzymatic reactions. The present work, as other works before (Holling, 1959; Arditi and Ginzburg, 1989; Garvie, 2007), clearly suggest that the usage of ordinary and partial differential equations can be extended more accurately to the realm of biology and ecology by appropriately modifying the law of mass action. Nevertheless, some limitations are not surmountable and must be utterly outlined. Firstly, the fact that the model parameters are changing as the dynamics unfolds, since the tumor morphology is evolving, specially when the tumor growth is considered. As shown in other works, tumors become disconnected during their interaction with the cell-mediated immune response for long times (López et al., 2017d). Secondly, the fact that the parameters change as a tumor becomes infiltrated. Therefore, the usage of stochastic discrete spatiotemporal models, such as cellular automata and agent-based models becomes indispensable to capture many of the complicated details of biological evolution and the dynamics of ecological and social systems.

